# Evaluation of somatic variant calling methods on high coverage tumour-only amplicon sequencing data in a clinical environment

**DOI:** 10.64898/2026.04.08.717310

**Authors:** Dhammapal Bharne, Daniel Gaston

## Abstract

One of the current workhorses of next-generation sequencing in clinical molecular diagnostics laboratories for profiling somatic mutations in tumours are amplicon-based targeted sequencing panels. Many open-source somatic variant callers are available; however, their use in clinical applications remains under explored. Therefore, we integrated outputs of six variant callers (FreeBayes, MuTect2, Pisces, Platypus, VarDict and VarScan) into a Snakemake pipeline and evaluated tumour-only data from the HD789 commercial reference standard sequenced in triplicate on three different sequencing runs using the Illumina AmpliSeq Focus panel on MiSeq and NextSeq 2000. A 1:4 dilution sample was sequenced for evaluating limits of variant detection. The called variants were analysed along depth, allele frequency, and other sequencing metrics. The variant callers were evaluated by their level of concordance and performance on known somatic variants. FreeBayes consistently called the largest number of somatic variants in each sample but also included more potential artifacts. Overall, FreeBayes, VarScan, MuTect2, and Pisces had the best performance on HD789 data.

## Introduction

A variety of Next-Generation Sequencing (NGS) techniques have been developed to detect genomic variants in human cancer genomes, including whole genome sequencing, whole exome sequencing and targeted sequencing. The targeted sequencing currently dominates clinical NGS applications in cancer due to cost, input requirements, and the level of required sequencing infrastructure [Chatterjee et al., 2017]. It can produce very high depths of coverage, allowing for the detection of somatic variants present at low variant allele frequencies, especially in highly heterogeneous tumors [Xu et al., 2017, Cohen et al., 2021].

Targeted Amplicon sequencing uses specific primers to amplify only the regions of interest. Many studies have suggested varying levels of concordance among variant callers due to a number of factors, including algorithm choice, varying default thresholds, and differing variant calling strategies [Liu, et al., 2013, Cornish et al., 2015, Kroigard et al., 2016, Bian et al., 2018, Li et al., 2019, de Schaetzen van Brienen et al., 2020]. However, the majority of comparisons have been in the context of whole genome sequencing. This study examines the performance of several widely-used variant callers on the kind of high-depth targeted sequencing data produced by widely used clinical assays. We evaluate six variant callers: FreeBayes [Garrison et al., 2012], MuTect2 [Cibulskis et al., 2013], Pisces [Dunn et al., 2019], Platypus [Rimmer et al., 2014], VarDict [Lai et al., 2016], and VarScan [Koboldt et al., 2012].

HD789 is a well-characterized, formalin-fixed, paraffin-embedded (FFPE) commercial reference standard with a mix of mutations, including abnormalities not tested for the sequencing strategy employed. Replicate sequencing data of HD789 on the Illumina AmpliSeq Focus DNA panel [Illumina, San Diego, CA, USA] including replicates sequenced with differing levels of input dilution were evaluated using each of the six variant callers to measure performance characteristics as well as the impact of factors such as depth and allele frequency.

## Material and Methods

### SSVCC

FreeBayes, MuTect2, Pisces, Platypus, VarDict and VarScan were integrated into a Snakemake workflow that we have called SSVCC. SSVCC checks the quality of FASTQ files using FASTQC and performs adapter and quality trimming using FASTP [Chen et al., 2018]. Then, reads are aligned to the reference genome using BWA-MEM2 [Md et al., 2019]. Mapped reads are assigned to read groups and coordinate sorted. A base quality score recalibration table is generated with GATK BaseRecalibrator [McKenna et al., 2010], excluding regions around known polymorphic sites obtained from dbSNP [Sherry et al., 2001] and Gnomad [Karczewski et al., 2020]. The recalibration table is used to apply a linear base quality recalibration model using the BaseRecalibrator tool [Van der Auwera et al., 2013]. Finally, variants are called using FreeBayes, MuTect2, Pisces, Platypus, VarDict, and VarScan.

SSVCC accepts a user defined configuration file. If the configuration file is not supplied, it will automatically generate a configuration file based on the other supplied arguments. For example, it uses the argument -vc for specifying variant calling through FreeBayes, MuTect2, Pisces, Platypus, VarDict, and/or VarScan by using the argument values *freebayes, mutect2, pisces, platypus, vardict* and *varscan* respectively. The -vc flag can take one value or a comma separated list of values for choosing specific variant callers or it can also take a value of *all* for selecting all of these variant callers. The other arguments it takes include -d for minimum depth of a variant, -af for minimum allele frequency threshold of a variant default to 0.01 among other parameters. The argument --add_flags is used to add specific parameters to variant callers. Allocations of number of processors, maximum memory and disk space are managed by its arguments -n, -ms and -ds respectively.

The variants called through the SSVCC workflow are decomposed and left-normalized. The called normalized variants are stored in both VCF and HDF5 file formats by default. SSVCC implements its various steps such as pre-processing, mapping, BQSR, and variant calling in conda environments for cross-platform compatibility.

Several accessory Python and R scripts for downstream analysis of the called variants are included. For example, h5_query.py is designed to extract variants that fall within specified regions from HDF5 files using a provided BED file; V_vs_C.py (variants by variant callers) and V_vs_S.py (variants by sample) are used to generate a table of variants called by each variant caller in all the samples and a table of variants called in each sample by all the variant callers respectively; Pisces_chr_sizes.py is used to generate an index file of chromosome sizes which is required for variant calling by Pisces; and VEP.py is designed to annotate variants with Ensembl VEP [McLaren et al., 2016]. The R scripts vioplot.R and UpSet_Plots.R create violin plots of depth of sequencing across variants and overlapping variant calls respectively.

### HD789

HD789 is a commercial reference standard of FFPE treated DNA with a set of well characterized variants including SNVs, Indels, and structural variants (SV) with allele frequencies in the range of 5 to 15 percent. Three independent sequencing replicates of HD789, _S1, _S2, _S3and a 1 to 4 dilution sample S-dil were prepared. The sequencing libraries were constructed using the Illumina AmpliSeq Focus panel and sequenced on a MiSeq and NextSeq 2000. The panel contains 269 amplicons across 47 genes of clinical significance in solid tumours: AKT1, ALK, APC, AR, BIRC2, BRAF, BRCA1, CCND1, CDK4, CDK6, CTNNB1, DCUN1D1, DDR2, EGFR, ERBB2, ERBB3, ERBB4, ESR1, FGFR1, FGFR2, FGFR3, FGFR4, GNA11, GNAQ, HRAS, IDH1,, DH2, JAK1, JAK2, JAK3, KIT, KRAS, MAP2K1, MAP2K2, MED12, MET, MTOR, MYC, MYCN, NF1, NRAS, PDGFRA, PIK3CA, RAF1, RET, ROS1, and SMO. The sequencing data obtained was then analysed using SSVCC with the hg38 human reference genome with a minimum depth of one and a variant allele frequency of at-least 0.0005. The mean depth of coverage across amplicons was analysed using Sambamba [Tarasov et al., 2015]. On-target variants were extracted from VCF files output from each variant caller. Distributions of number and types of variants were determined for each variant caller. The variants were analysed using depth, allele frequency, filter parameters, and other sequencing metrics. Annotations and significance of the variants was determined by evaluating through VEP [McLaren et al. 2016]. Concordance among the variant callers was estimated by considering the variants called by at least two different variant callers in any of the replicates and the diluted sample. Further, the performance of the variant callers was ascertained by whether variant callers did or did not successfully identify known variants from the HD789 sample that fell within the target regions of the AmpliSeq Focus sequencing panel.

## Results

### Survey of variant callers for tumour-only targeted somatic sequencing applications

A survey of the literature for variant callers that would identify small variants (SNVs, MNVs, and small indels) on tumour-only data in a calling mode suitable for identifying somatic variants was performed and initially identified none different callers: 16GT [Luo et al., 2017], FreeBayes [Garrison et al., 2012], LoFreq [Wilm et al., 2012], MuTect2 [Cibulskis et al., 2013], Pisces [Dunn et al., 2019], Platypus [Rimmer et al., 2014], VarDict [Lai et al., 2016], VarScan [Koboldt et al., 2012] and DRAGEN [Behera et al., 2025]. However, 16GT requires the use of SOAP-dp aligner [Luo et al., 2013] and LoFreq proved difficult in terms of computational memory resources on the high-depth, targeted sequencing evaluated here and after initial testing both were dropped from the analysis. Because DRAGEN needs specialized hardware or Illumina specific cloud environments for analysis it was also removed from further consideration. The remaining variant callers were considered feasible for routine clinical applications and deployment on reasonable hardware in a standard Linux environment, and integrated into the SSVCC variant calling framework. A directed acyclic graph (DAG) of the SSVCC workflow is shown in Figure 1.

**Figure 1.**
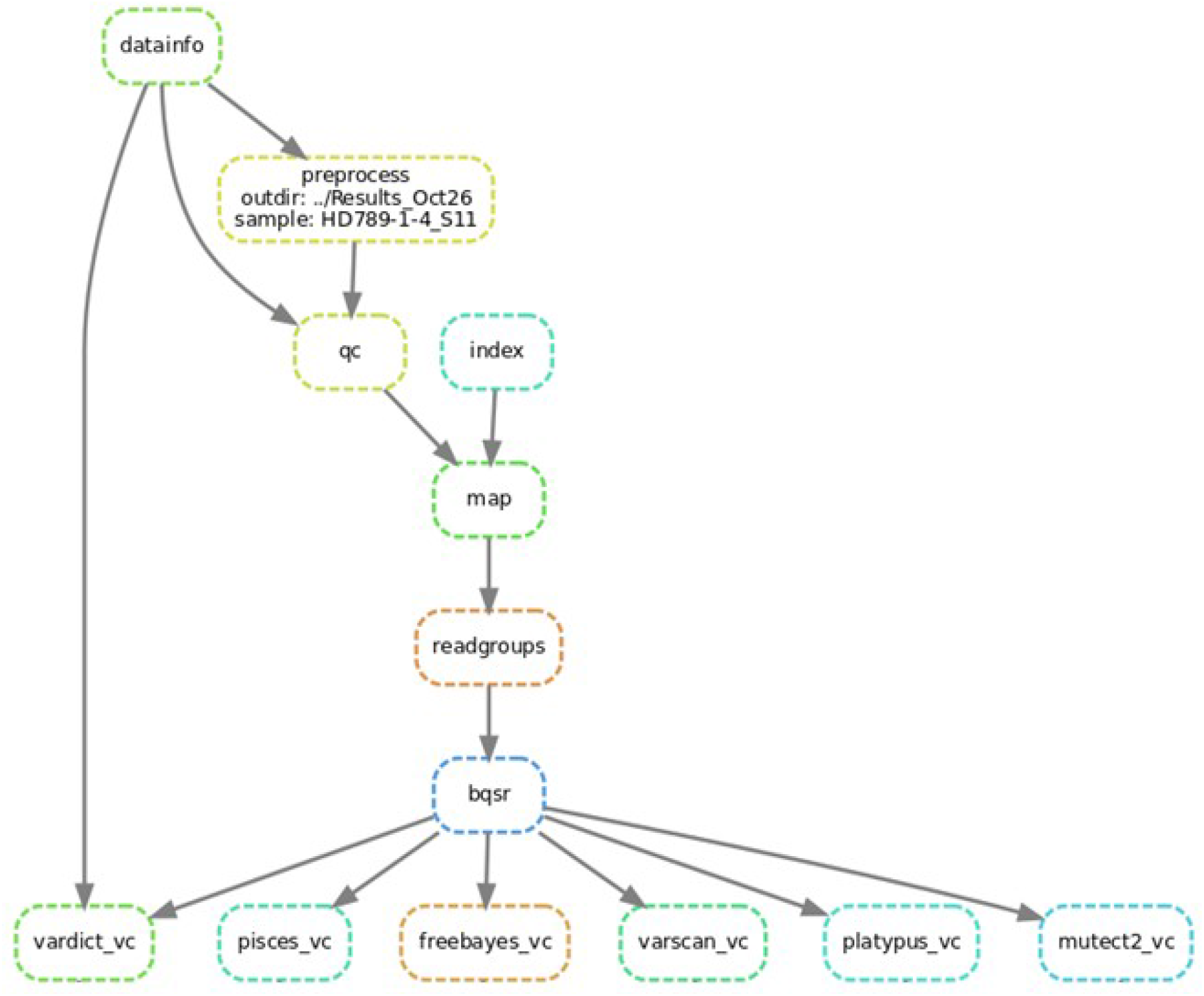
Directed acyclic graph (DAG) of data processing and variant calling by SSVCC. It includes pre-processing, mapping, and recalibration steps followed by variant calling with FreeBayes, MuTect2, Pisces, Platypus, VarDict, and VarScan.

### Amplicons coverage

Mean depth of coverage of the Focus panel amplicons were calculated using Sambamba [Tarasov et al., 2015]. The mean depth was lowest in HD789_S2 and the highest in HD789_S3. The lowest mean depth amplicons in HD789_S2 as well as in HD789_S3 were found in FGFR3, while in HD789_S11 as well as in HD789_S-dil it were in FGFR4. The highest mean depth amplicon was MYC in all of these samples. The mean depth of coverage of all amplicons on the panel and across all samples is shown in Table 1 (Table S1). More than 90 percent of the amplicons in the targeted regions in HD789_S3 were covered at a depth of at least 2000X, nearly 80 percent in the 1:4 dilution sample, and close to 50 percent in the other HD789 replicates. However, at 1000x, the percent of amplicons in the targeted regions was above 90 in all except HD789_S2 where it was 84 percent.

**Table 1.** A table of filters, depth and allele frequency ranges of the on-target variants.

| **Variant Callers** | **On-target Variants** | **Filters Field Values** | **Depth Range** | **Allele Frequency Range** |
| --- | --- | --- | --- | --- |
| FreeBayes | 4069 | PASS | 234 - 26187 | 0.00500 - 1 |
| MuTect2 | 622 | PASS, base_qual, clustered_events, fragment, germline, haplotype, multiallelic, orientation, slippage, strand_bias, weak_evidence | 19 - 38913 | 0.00067 - 0.999 |
| Pisces | 419 | PASS, R5x9, SB, q30 | 312 - 21998 | 0.01410 - 1 |
| Platypus | 164 | PASS, QD, SC, alleleBias, badReads, strandBias | 157 - 6141 | 0.01810 - 0.99966 |
| VarDict | 847 | LongMSI, MSI12, PASS, Q10, p8, pSTD | 83 - 24233 | 0.00510 - 1 |
| VarScan | 442 | PASS | 80 - 10840 | 0.00500 - 0.999 |

### On-target variants

In all 2655 unique variants were called across all replicates. The distribution of types of on-target variants identified by different variant callers across all replicates is shown in Figure 2.

**Figure 2.**
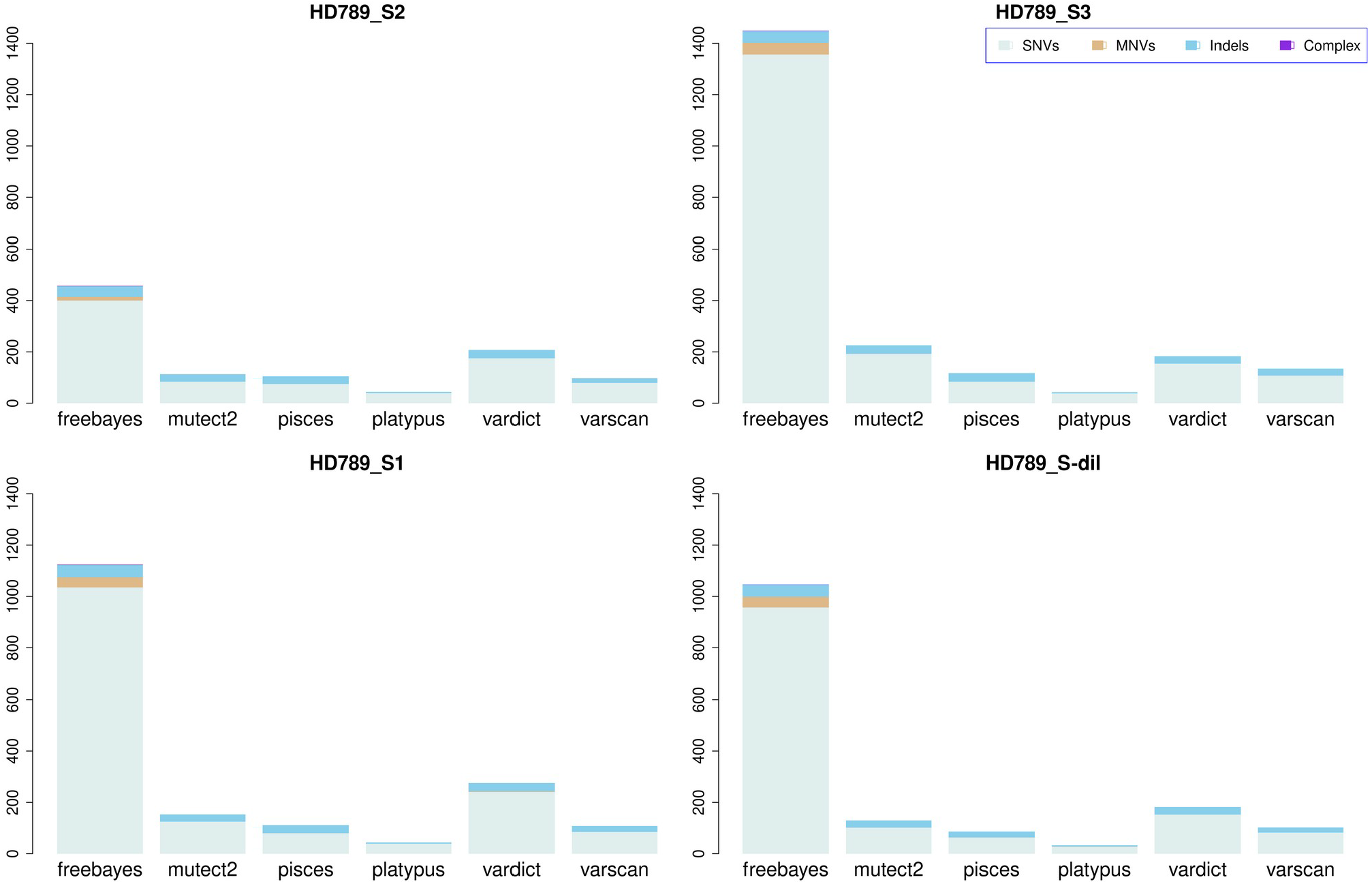
Breakdown of on-target variants by type called by each of the tested variant callers in HD789 replicates (HD789_S1, HD789_S2, HD789_S3) and the diluted sample (HD789_S-dil). SNVs, MNVs, indels, and complex variants are colour coded in stacked bar graphs.

FreeBayes makes the highest number of variant calls (2433) while Platypus the lowest (55). Despite being replicates of the same commercial control sample there were differences in the number of per-sample variant calls even within the same variant caller with the highest seen in HD789_S3 (1570) and the lowest in HD789_S2 (483). In all the samples, FreeBayes makes the highest number of variant calls and Platypus the lowest. 140 MNVs were called by FreeBayes in each of the replicates and the dilution sample while two MNVs, chr5_177096558_CGTC_TGTA and chr5_177096559_GTC_ATA, were called by VarDict only in HD789_S1 and one MNV, chr5_177090796_CCTCG_TCTCA, was called by Platypus only in the diluted sample. The other variant callers did not call any MNVs in any of the samples. Several more complex indel/substitution variants were called only by FreeBayes in one or more replicates (chr12_25233126_CCAC_A in HD789_S-dil (depth=2722, allele frequency=0.24779), HD789_S1 (depth=2203, allele frequency=0.07616) and HD789_S3 (depth=3581, allele frequency=0.04527) and chr4_54663033_T_CA in HD789_S2 (depth=234, allele frequency=0.02597). These findings suggested FreeBayes may be able to identify more complex variants but also is likely to produce an elevated rate of false positive artefactual calls.

### Variant depth of coverage and variant allele frequency

The distribution of depth of coverage of variants called by variant callers in each of the replicates is shown in Figure 3. The depth of coverage at called variants was found in the range of 19 to 38913. The distribution of allele frequency of the variants versus depth is shown in Figure 4. The allele frequency of the variants was found in the range of 0.00067 to 1 with the majority of variants, regardless of depth of coverage, falling below 0.2.

**Figure 3.**
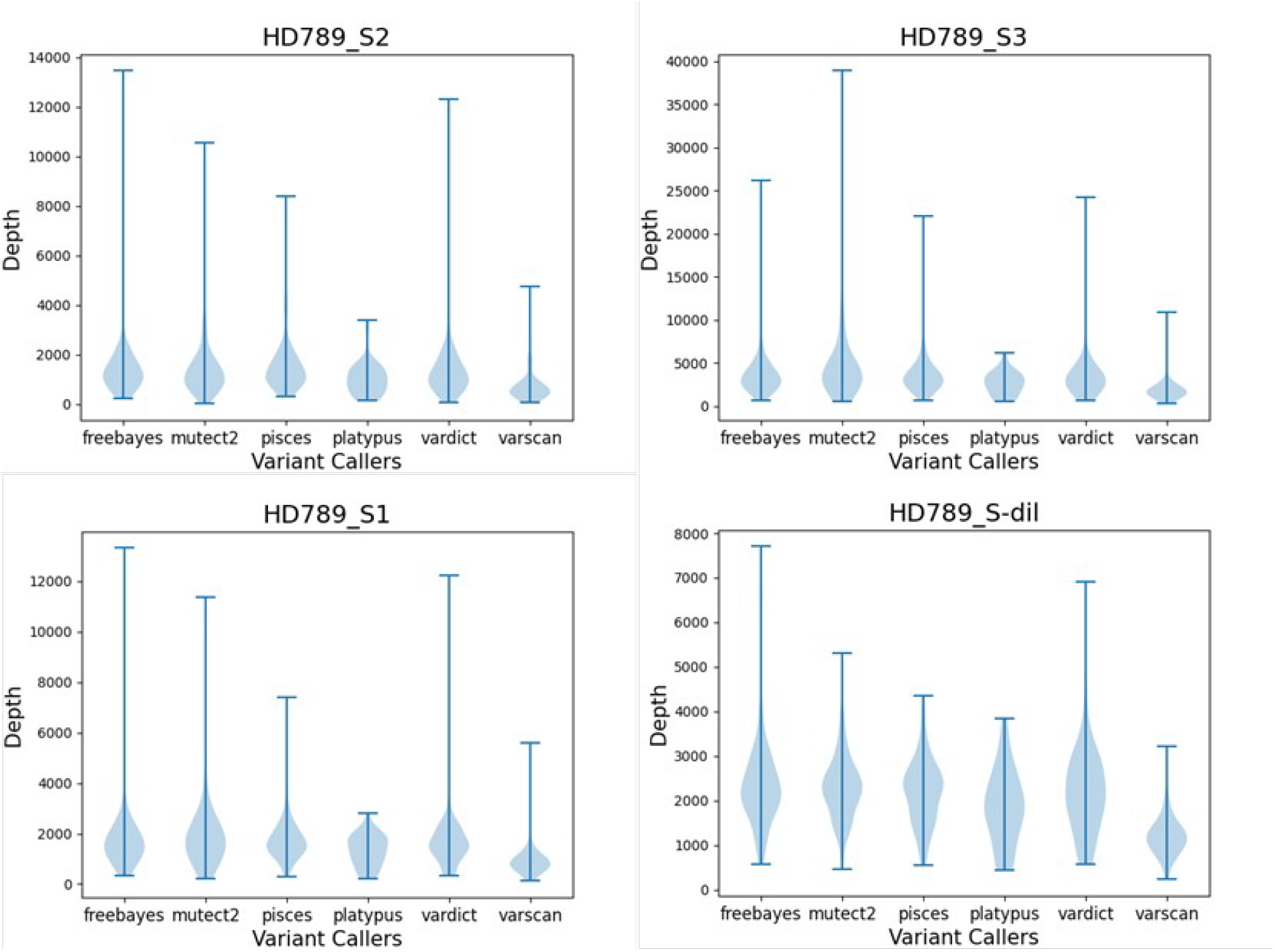
The distribution of depth of variants called by different variant callers.

**Figure 4.**
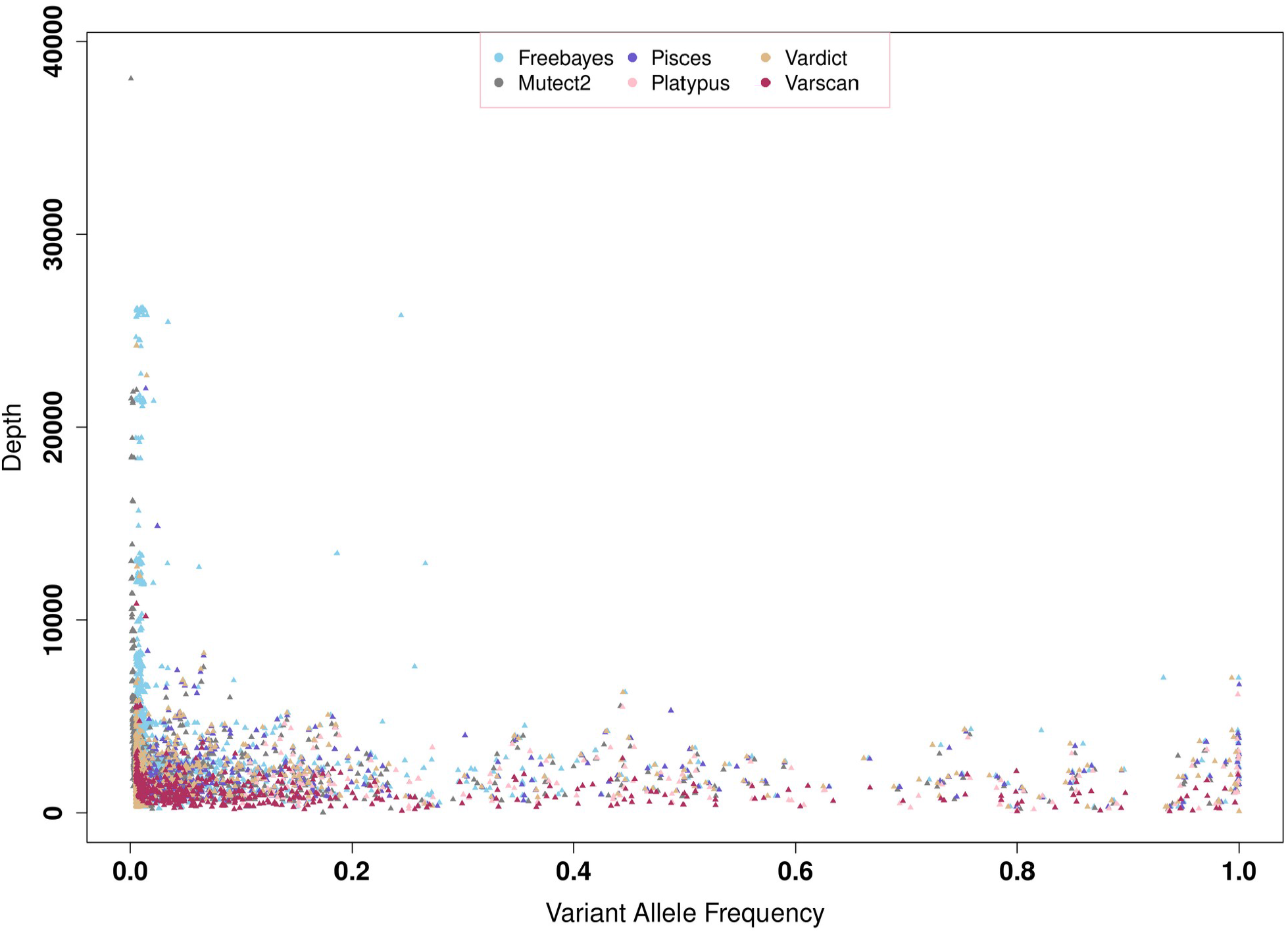
Scatterplot of allele frequency versus depth of the raw, unfiltered on-target variants, coloured by variant caller.

### Variant filtering

Filtering criteria and the resulting filter field flags differ between variant callers. For example, MuTect2 uses clustered events, orientation bias, slippage, germline evidence among others while Platypus uses allele bias, strand bias, etc. A list of filters applied by the variant callers on the on-target variants is shown in Table 1. The table also show the maximum and minimum depths and allele frequencies of the on-target variants called by each variant caller.

The variants that did not trigger any of the filters set by a variant caller are marked as PASS. Mutect2, Pisces, Playtpus, and VarDict were the only variant callers running filters here that were triggered in any variants called. Filtering primarily on the filter field of a VCF for those marked as PASS is a common filtering strategy in variant analysis. The number of PASS variants, on-target variants is shown in the following Figure 5. While FreeBayes and VarScan only contained variants marked as PASS this is primarily due to a lack of specific filters running under default parameters. The distributions of depth of coverage and allele frequency of the PASS and non-PASS variants is shown in Figure 6.

**Figure 5.**
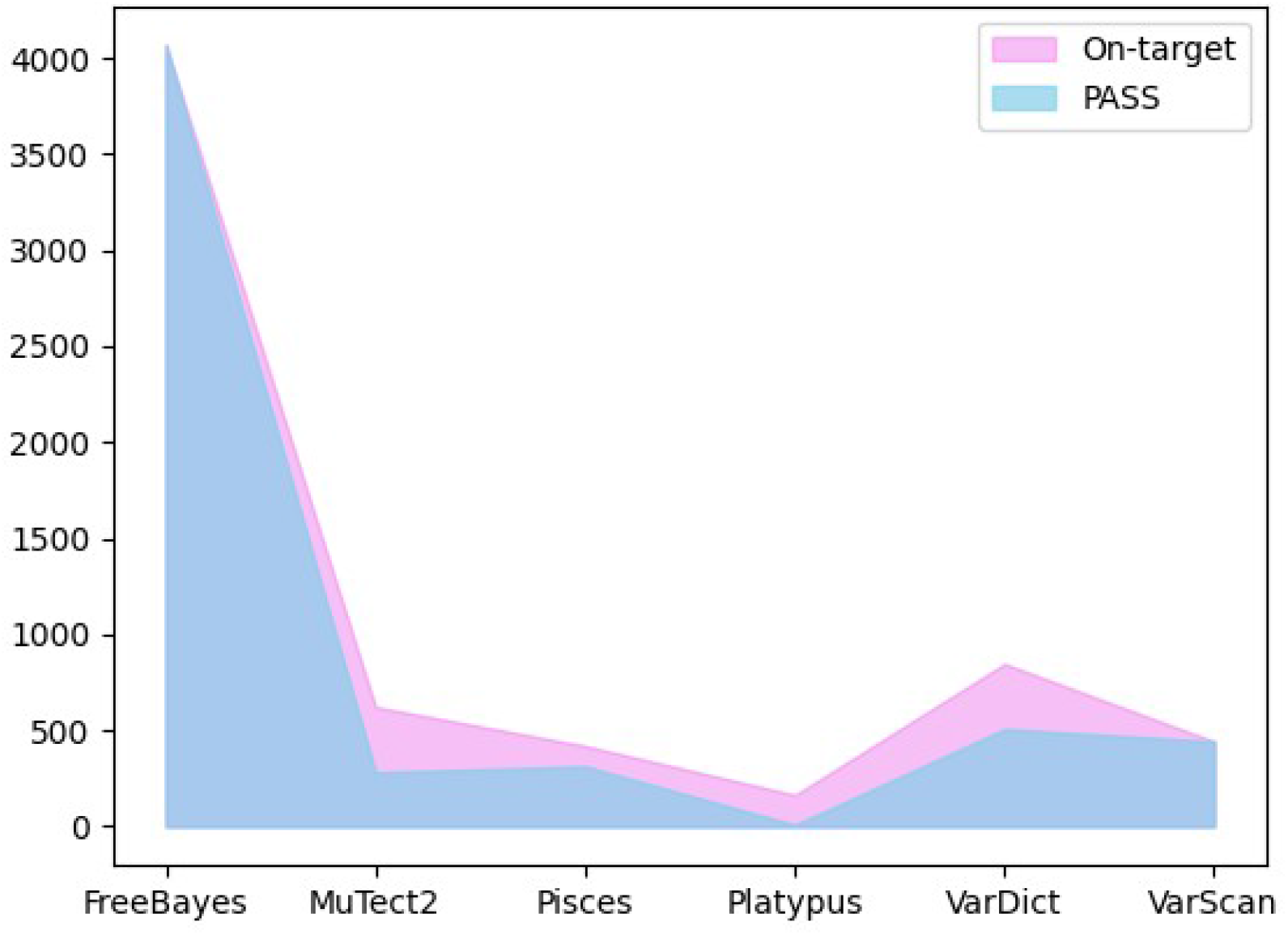
The distribution of on-target and passed variants.

**Figure 6.**
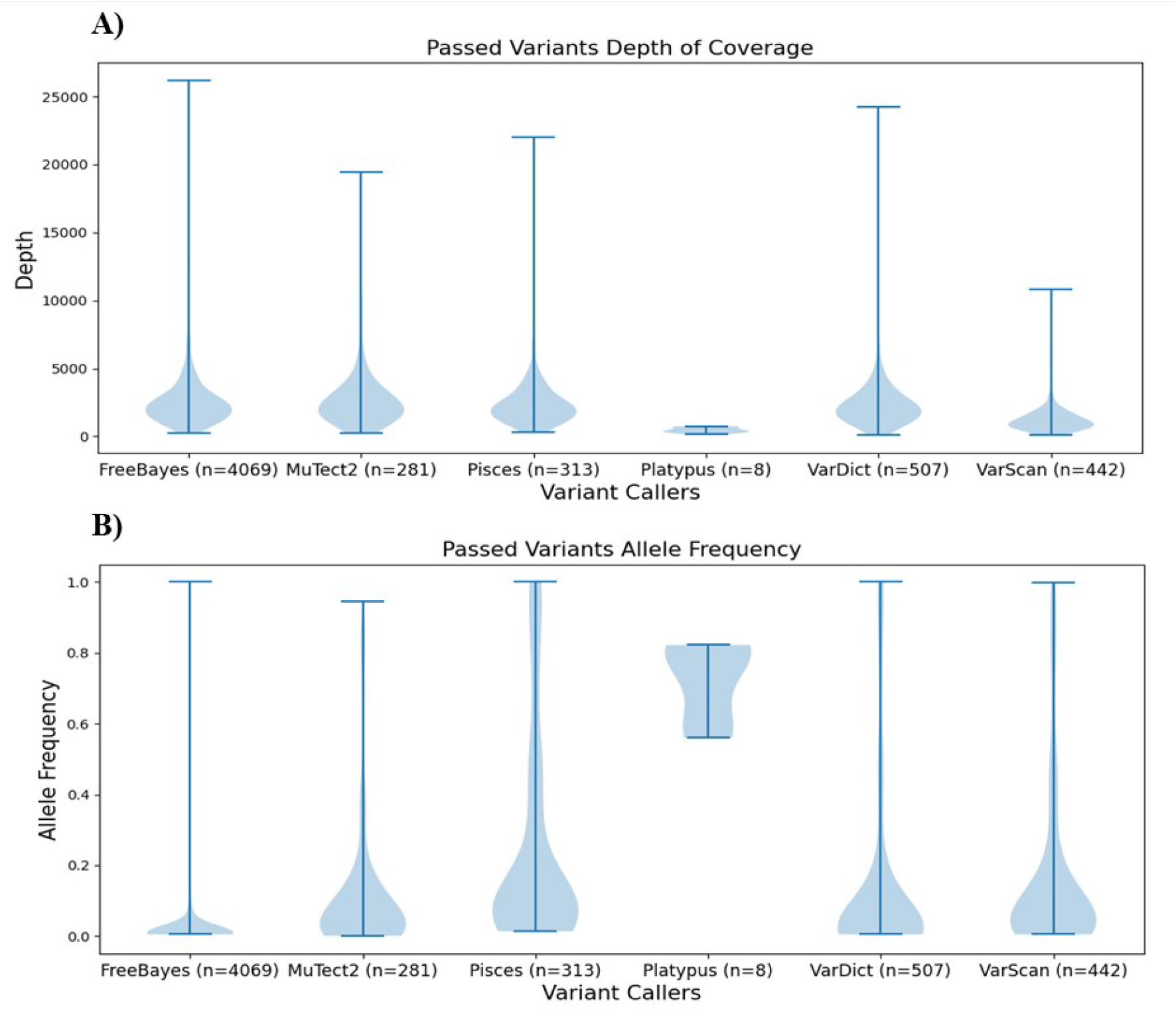
The distribution of depth A) and allele frequency B) of passed variants

### Variant consequences

VEP annotation results indicated that, of the 2655, 788 variants were known while the renaming 1875 were novel. It also summarized the on-target variants to overlap with 155 genes, 1310 transcripts and 77 regulatory features [McLaren et al., 2016]. Most of the on-target variants were modifiers originating from introns, upstream or downstream region of the genes. There were many coding sequence variants such as missense, inframe insertion or inframe deletion variants. These coding sequence variants were observed to have moderate effects with change in the amino acid sequence without changing the frame of the transcript. There were some variants with low consequences such as synonymous, start or stop retained variants. These variants were observed to change codon sequence without changing the amino acid. A few high effect variants were observed which consisted of frameshift, stop-gain or loss, and splice acceptor or donor variants. These variants were observed to interfere with splicing, transcription and translation processes causing significant effects in the downstream signalling and cellular pathways. Further, some of the low to moderate and modifier consequence variants were clinically known with benign or pathogenic significance, however, all the high effects variants were clinically known but with uncertainty significance. The following Table 2 summarizes high-impact variants detected from VEP [McLaren et al. 2016] by each variant caller across replicates along with their variant allele frequency.

**Table 2.**
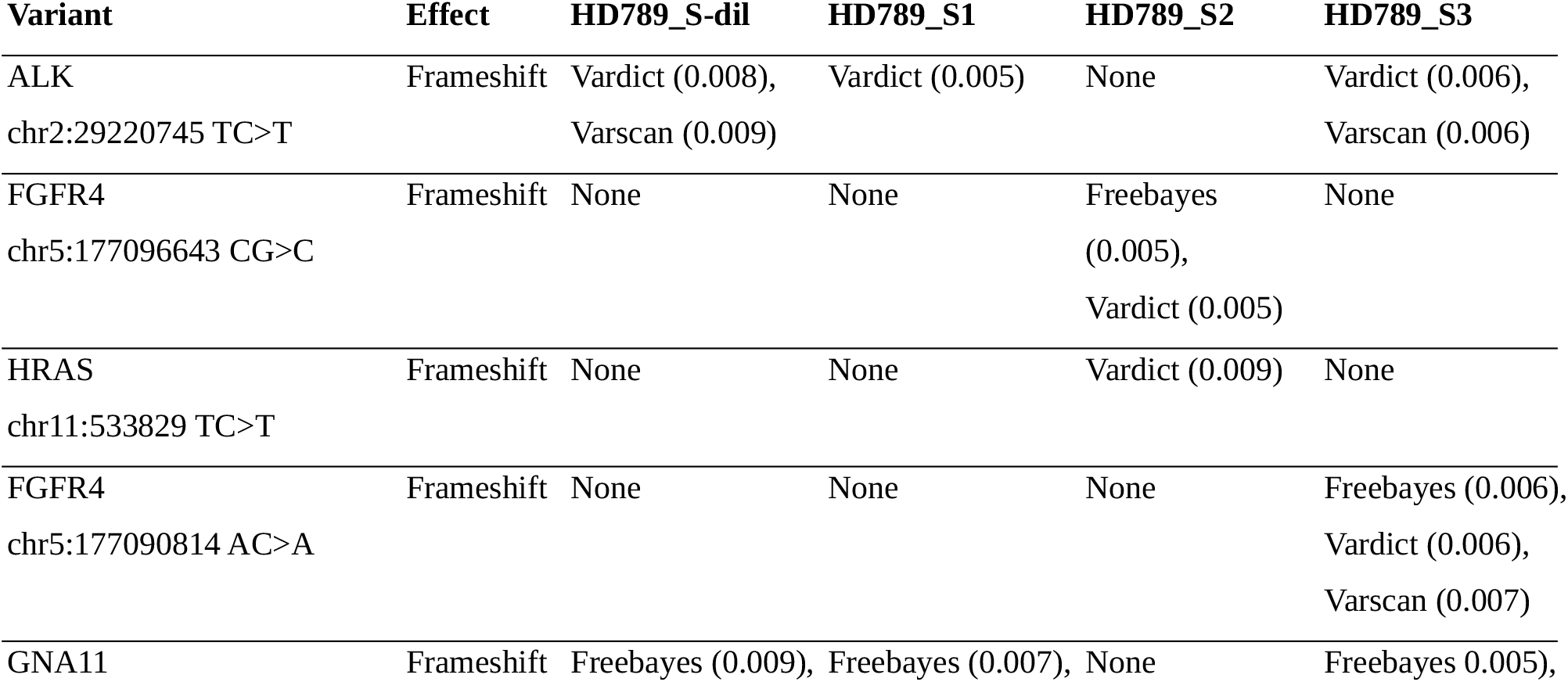
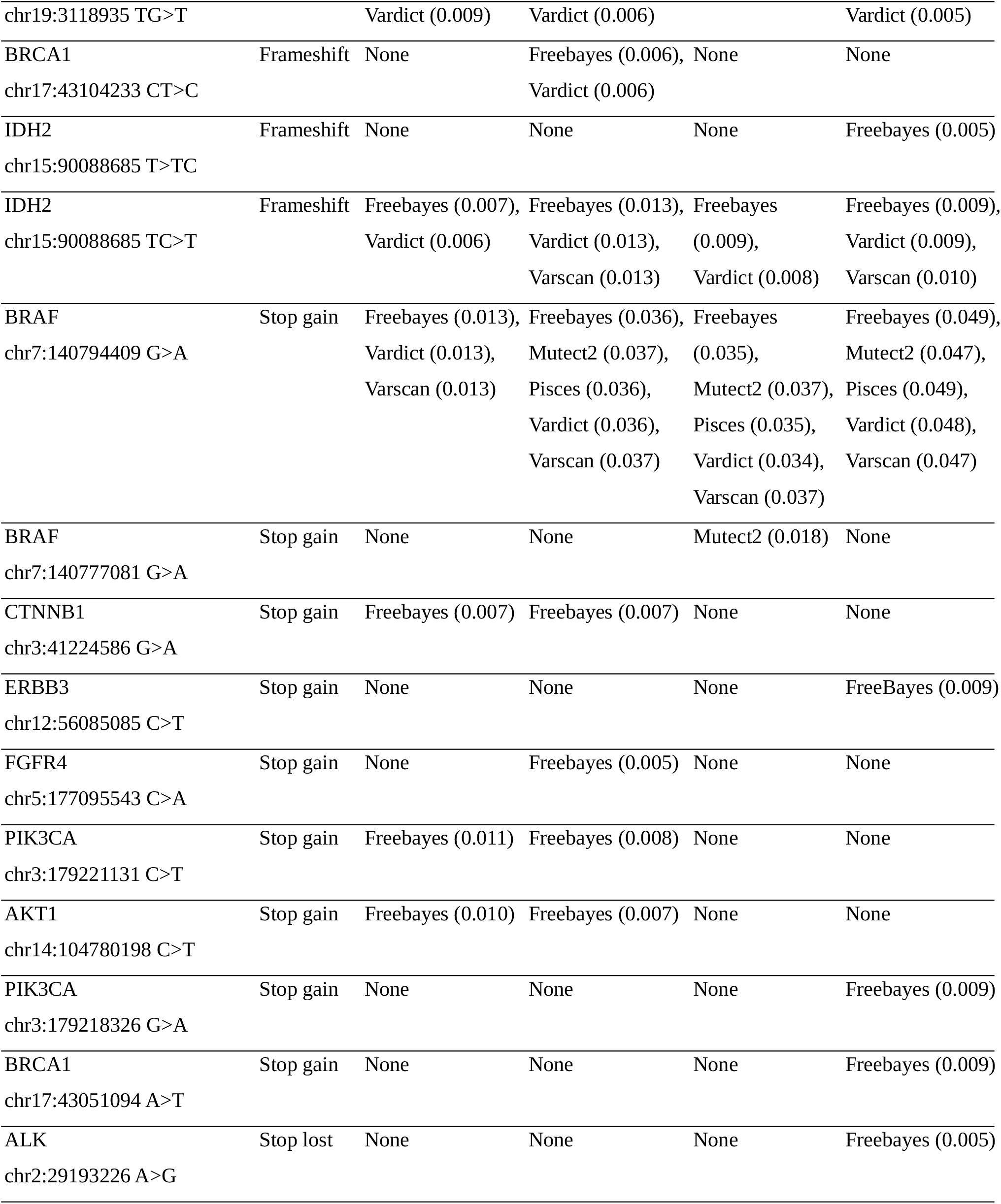
Summary of high impact variants (frameshift, top gain, or stop loss) from the on-target variants called. The table also includes sample wise variant callers calling these high effect variants. Allele frequency of the variants called by the variant callers are enclosed in the round brackets.

### Variant callers’ concordance

Of the total on-target variants, 609 were called by at-least two different variant callers in one or more replicates, including the dilution replicate, suggesting good concordance among the variant callers tested. The following Figure 7 shows the number of variants called by different combinations of variant callers in the dilution replicate as an example. FreeBayes and VarDict call a much higher number of concordant variants than any other combination of the variant callers. However, these two variant callers (along with VarScan) call the highest number of variants overall, with FreeBayes much higher than any other caller. The next two most concordant combinations of callers were all callers, with or without Platypus (which calls the fewest variants overall).

**Figure 7.**
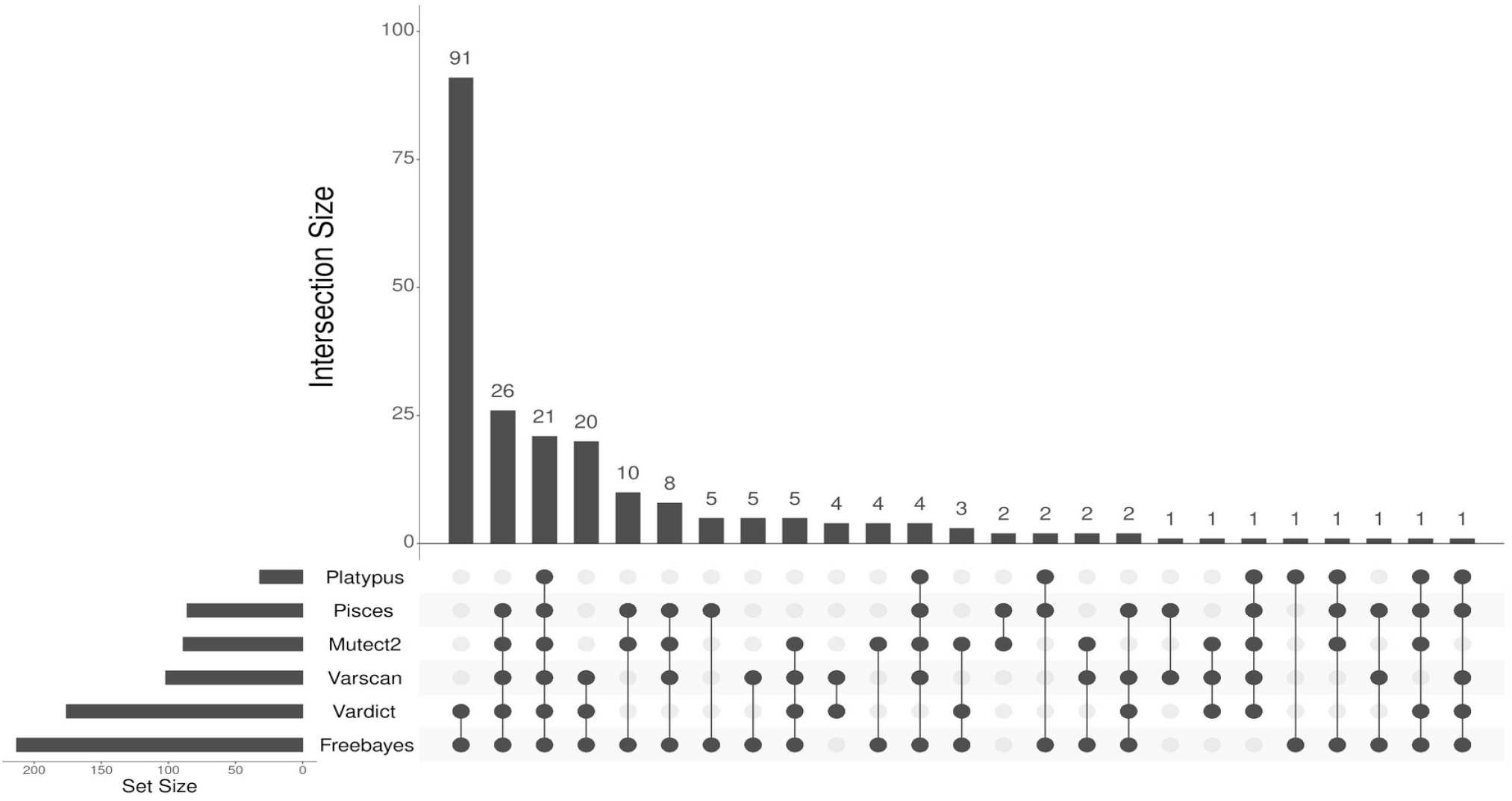
Upset plot demonstrating concordance between on-target variants called by at least two different variant callers. Intersection size (y-axis) is the number of variants called by all variant callers shown below with dots connected by lines. Set size is the total number of on-target variants called by the variant caller.

### Variant callers’ performance on known variants in HD789

The performance of the variant callers was evaluated on known variants in HD789 that were 1) covered by the Illumina AmpliSeq Focus panel and 2) of a variant type detectable by targeted sequencing and the small variant callers evaluated here. The present study considered fifteen mutations: 3 SNVs of PIK3CA and a single SNV each of DDR2, CTNNB1, EGFR, BRAF, KRAS, AKT1, MAP2K1 and GNA11 genes. It also contained a deletion mutation of each of CTNNB1 and EGFR, a duplication mutation of EGFR and a frameshift mutation of FGFR3. The following Table 3 lists all these mutations in HGVS format.

**Table 3.**
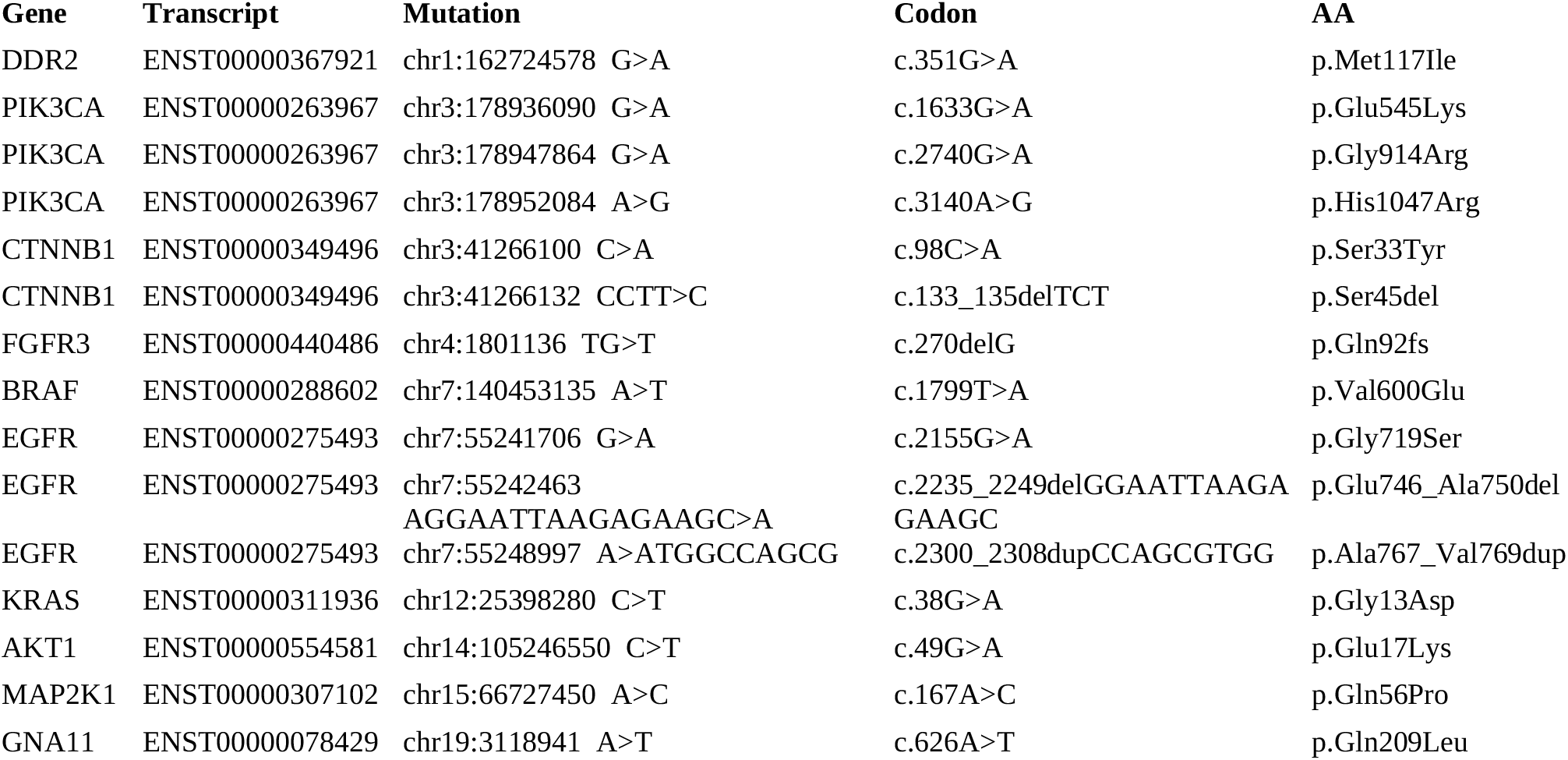
List of known mutations in HD789.

**Table 4.**
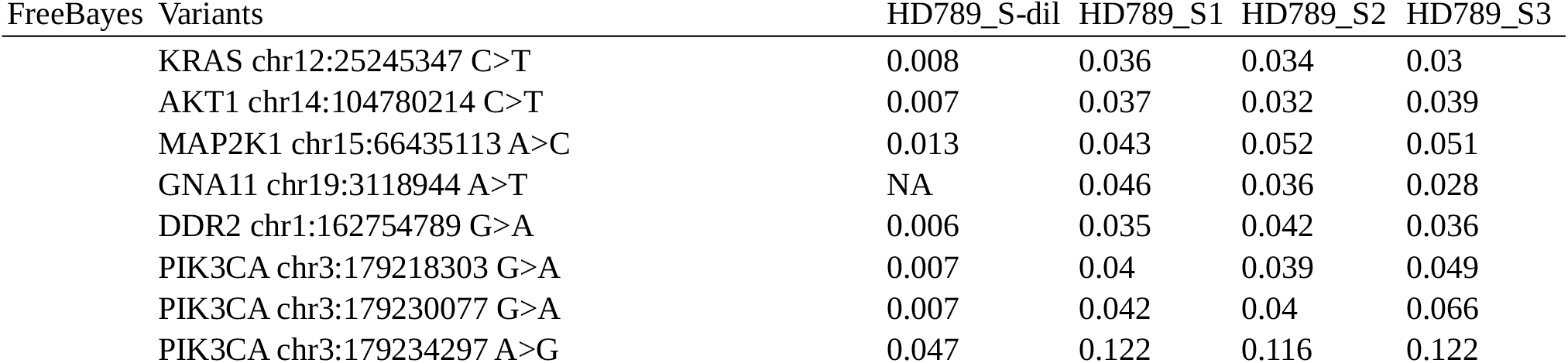
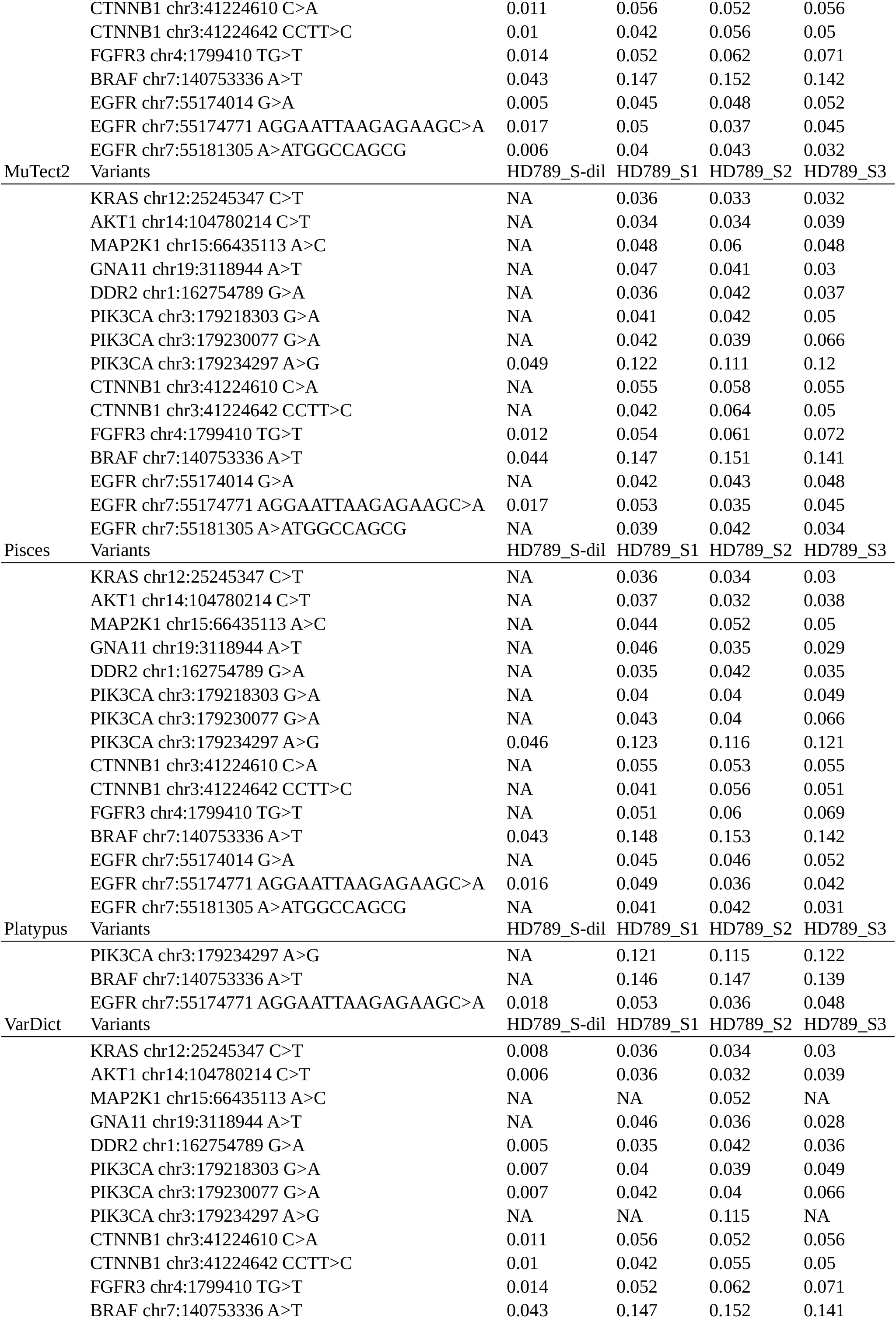
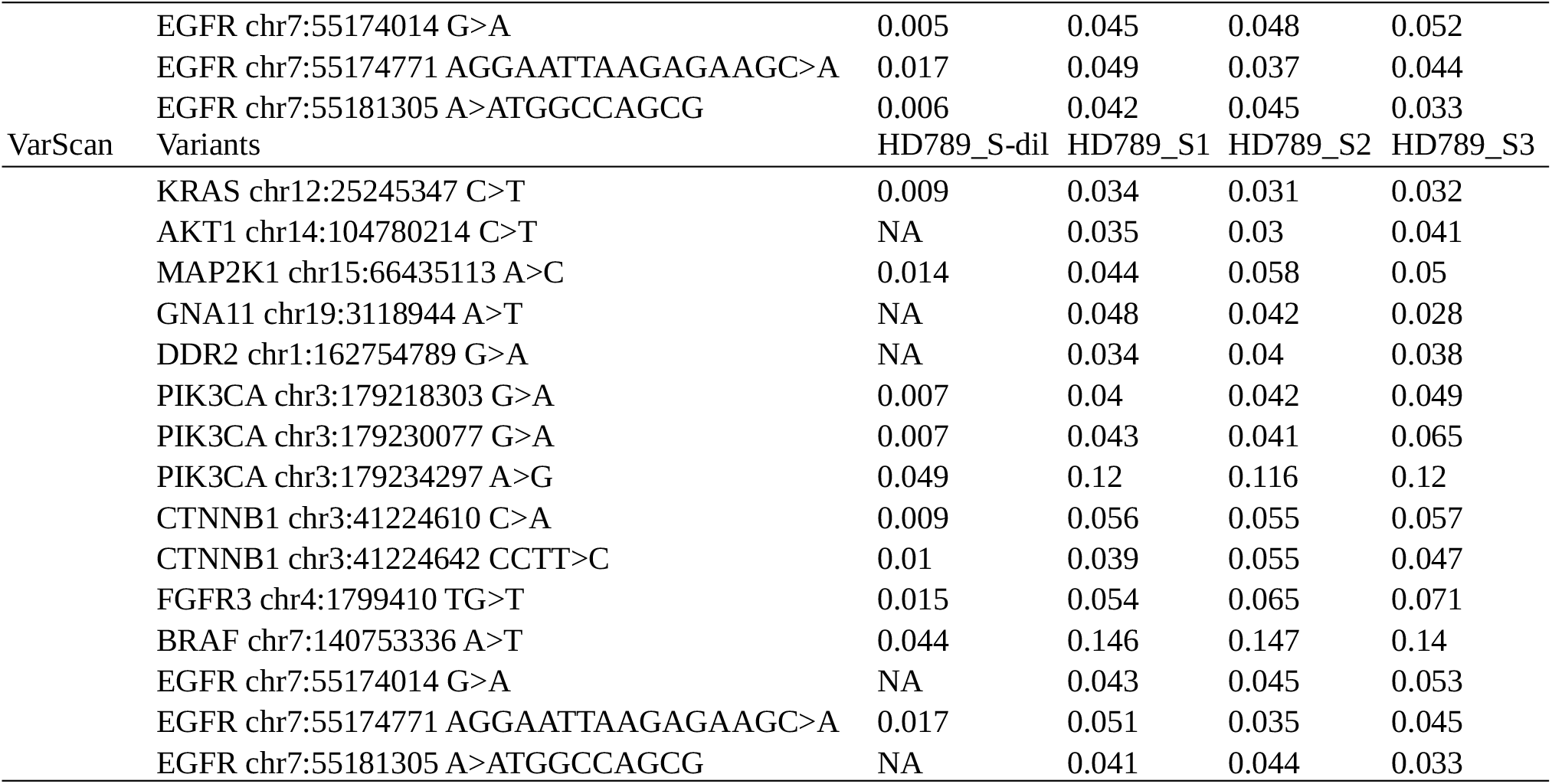
Variant allele frequencies of known variants in HD789 in each replicate by variant caller.

Figure 8 shows the performance of each variant caller for each of these mutations. FreeBayes detected all mutations across all replicates with the exception of GNA11 in the input DNA dilution replicate. Platypus had the worst performance, successfully calling only three of the known mutations in the standard input replicates and only one in the dilution replicate. It called the EGFR deletion mutation (EGFR p.Glu746_Ala750del), PIK3CA (PIK3CA p.His1047Arg), and BRAF (BRAF p.Val600Glu) in the standard replicates but missed these PIK3CA and BRAF mutations in the dilution replicate. FreeBayes successfully identifies the greatest number of known well characterized variants in the HD789 commercial control sample, but also has the highest overall number of variant calls, many of which are low-quality or artefactual. Mutect2, Pisces, VarDict and VarScan have comparable performance to one another, with slight variations in calling. The worst performance was generally on the diluted input sample, a factor which needs to be considered carefully when interpreting outputs from such samples.

**Figure 8.**
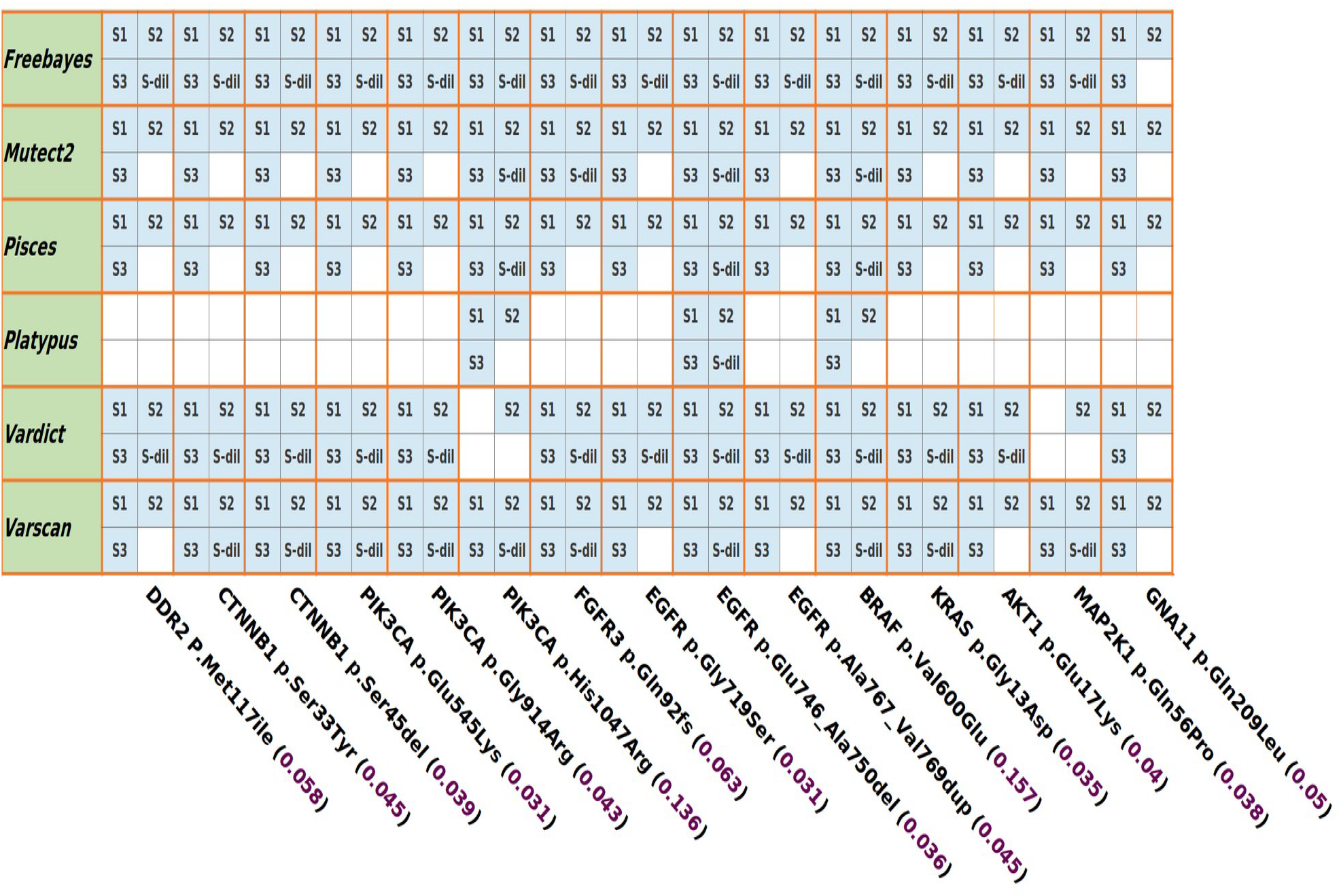
Performance of each variant caller across all three replicates and the 1:4 dilution sample of well characterized variants in HD789 with their known variant allele frequencies. If the variant was called by a variant caller in a particular replicate or the dilution sample it is represented by a filled in light blue coloured box. If a variant was not called it is represented by an empty box. S1, S2, S3 are HD789_S1, HD789_S2, HD789_S3 replicates respectively and S-dil is HD789_S-dil diluted sample.

A table of variant allele frequencies of these mutations as produced by the individual caller in the individual replicate is shown in the Table 3 along with the ddPCR verified variant allele frequency of the sample from the manufacturer.

## Discussion

Despite many advancements in tumour analysis and variant calling, there remain a paucity of open-source variant callers capable of running in a tumour-only somatic sequencing mode, particularly with high-depth sequencing data like that seen in targeted sequencing applications. To date comparisons of variant calling performance have largely focused on the whole exome or whole genome sequencing cases and typically with tumour-normal matched sequencing data. While these applications are important, and targeted sequencing applications are largely focused on the detection of known driver mutations, high-depth of coverage allows for the detection of low variant allele-frequency variants in the context of tumour-heterogeneity and sub-clonal evolution which can be critical for proper diagnosis, prognosis, or therapeutic selection in clinical sequencing cases. This work provides some initial insights of performance of a variety of open-source variant callers designed to operate in this mode on a FFPE commercial control sequenced with a targeted sequencing panel that is widely used in clinical sequencing applications and under typical clinical laboratory sequencing conditions.

The best performing variant callers under default parameters produced call sets enriched for variants called with lower depths of coverage and lower variant allele frequencies. While some of these variants will be “real”, many will also be sequencing or analytical variants, highlighting the importance of careful filtering and downstream analysis strategies. A good concordance was observed among the variant callers for on-target variants; however, only FreeBayes, MuTect2, Pisces, and VarScan called most of the known clinically significant variants across all replicates with the exception of the dilution replicate. Employing multiple variant callers with a concordance metric along with careful filtering and flexible depth/variant allele frequency thresholds will produce the most reliable call set. Different metrics/thresholds may need to be employed for known, well established, clinically actionable variants compared to novel variants especially with knowledge of input quantity/quality metrics for sequencing libraries.

